# Exploring Clinical Class 1 Integrons as Valuable Targets for the Re-sensitization of Multidrug Resistant Pathogenic Bacteria Using CRISPR-Cas

**DOI:** 10.1101/2025.03.28.645910

**Authors:** Anna Berggreen Skovmand, Nicolai Juel Paaske, Arancha Peñil-Celis, Amalie Elisabeth Schønemann, Lærke Lund Hansen, Witold Kot, M. Pilar Garcillán-Barcia, Tue Kjærgaard Nielsen

## Abstract

The increasing prevalence of bacteria resistant to many or all types of antibiotics poses a major health crisis. Novel classes of antibiotics are only slowly being developed and alternative strategies are needed to tackle the issue. Mobile genetic elements and class 1 integrons are important facilitators for antibiotic resistance genes, with the latter being highly conserved in human pathogens. The growing prevalence of multidrug-resistant bacteria and the paucity in the development of new antibiotics underscore the urgent need for innovative approaches in the treatment of pathogens. Among these, CRISPR-Cas nucleases can be used to cleave acquired resistance genes, leading to either plasmid curing or cell death if the target is on a chromosome. In this study, we investigate the feasibility of using class 1 integrons as a target for Cas9-based cleavage leading to re-sensitizing antibiotic-resistant bacteria. We analyze the conserved and widespread integrase gene *intI1* and conclude that it is a suitable target for Cas-based re-sensitization due to its high sequence conservation and its occurrence largely limited to human pathogens, alleviating the risk of targeting benign bacteria. We developed a broad host range conjugative plasmid encoding a class 1 integron-targeting Cas9 system that leads to removal of resistance plasmids in target bacteria with subsequent re-sensitization towards antibiotics. We find that 290 distinct ARGs co-occur on *int1*-harboring plasmids, showing the potential for re-sensitization towards a very broad range of antibiotics.

## Introduction

The spread of antibiotic-resistant pathogenic bacteria is a major threat to human health and a source of global concern^1^. Despite this, the discovery of new antibiotics has stalled, increasing the urgency to develop new approaches to combat drug resistance^2,3^. The utilization of CRISPR-Cas (clustered regularly interspaced short palindromic repeats; CRISPR-associated protein) systems for the precise targeting of resistance elements in bacterial populations presents a promising approach for new antimicrobials^4–8^. Antibiotic resistance can be mediated by chromosomal or plasmid-borne genes; however, acquired multidrug resistance (MDR) is mainly attributed to horizontal gene transfer (HGT) of antibiotic resistance genes (ARGs) via mobile genetic elements (MGEs) such as plasmids, transposons, and integrons^9–15^. Antibiotic resistance is often conferred by overexpression of ARGs^16^, where strong promoters within insertion sequence (IS) elements and integrons play a key role in this overexpression^14,17,18^.

Integrons are genetic elements that acquire, rearrange, and express gene cassettes^14^. They can be assigned to five different classes^10,14^, of which class 1 integrons have the greatest clinical significance^10,11^, exhibiting detection rates of up to 90% among Gram-negative pathogens from clinical samples^14,19^. Class 1 integrons share a common ancestor, as evidenced by their highly conserved *intI1* gene. They likely entered commensal bacteria through food or water, with the human gut microbiota as the initial colonization site^14,20,21^. Class 1 integrons consist of core elements: an integrase gene (*intI1*), which catalyzes site-specific recombination between the recombination site (*attC*) of a gene cassette and the integron attachment site (*attI*), and a common promoter (Pc) that enables transcription of gene cassettes^14,20^. Class 1 integrons carry nearly all known classes of acquired ARGs as gene cassettes with more being discovered continuously^21,22^. As such, they are crucial in the evolution of human pathogenic bacteria by facilitating the spread of ARGs^10,11,14,17^. While the majority of environmental integrons (not class 1) are found on bacterial chromosomes and carry cassettes encoding a variety of functions, clinical class 1 integrons are associated with MGEs such as plasmids and transposons^23,24^. The class 1 integron integrase activity is enhanced in response to internal SOS signals and DNA damage in its host^21^ and is further up-regulated during conjugative plasmid transfer^25^. Analysis of environmental integrases from metagenomic datasets shows low nucleotide sequence conservation^26^. In contrast, *intI1* genes in clinical pathogens are essentially identical^14^, which, combined with their mobility, abundance, and global distribution, makes class 1 integrons relevant targets for Cas-based antimicrobials^10^.

The genome-editing tool CRISPR-Cas system has numerous applications, including the development of novel antimicrobials^27^. CRISPR-Cas-based antimicrobial strategies employ two approaches: inducing bacterial cell death or eliminating/silencing ARGs^5,27–29^. Curing ARG-plasmids or disrupting ARGs on chromosomes can re-sensitize bacteria to antibiotics or kill the target cell, respectively. This shows a strong promise for CRISPR-Cas based antimicrobials^4,27,28^. Previous studies on CRISPR-Cas based antimicrobials have targeted specific ARGs and resistance mechanisms, such as genes encoding extended-spectrum beta-lactamases (ESBL)^4,30,31^. However, targeting a specific ARG with CRISPR-Cas antimicrobials has several drawbacks. In the example of ESBLs, there are multiple classes and sequence variants, such as *blaSHV, blaOXA, blaNDM, blaTEM, blaKPC, blaPER*, and others. The TEM-type ESBLs alone exhibit 250 different variants (i.e., sub-terms; CARD database^32^).

It is hypothesized that the evolution of beta-lactamases predates the clinical introduction of beta-lactam agents by more than two billion years^33^. These ancient enzymes have diversified extensively in both sequence and function, particularly through coevolution with biosynthetic pathways^34^. As a result, they represent a highly heterogeneous group of enzymes, with variants that can be traced back to different evolutionary origins^35^. Furthermore, beta-lactamases play a role in general cellular maintenance^36^, which could lead to the unintended killing of non-target bacteria by Cas-based antimicrobials. Each class of beta-lactamase has disseminated to a diverse array of bacterial species, and orthologous genes display significant sequence diversity, with sequence similarity as low as 20% observed between members of each class^33,34^.

One single ARG target sequence for CRISPR-Cas re-sensitization will therefore be of limited value. Acquired ARGs are distributed across phylogenetically unrelated bacteria due to HGT^37^. This poses another challenge to the development of a universal system or assay that can reach all ARG/bacteria combinations. In contrast, the class 1 integron-integrase gene (*intI1*) derived from clinical isolates is highly conserved, with nearly identical DNA sequences. This suggests a single common origin for all class 1 integrons currently identified in clinical pathogens^14,20^.

We argue that class 1 integrons offer an optimal target for re-sensitization to combat MDR pathogens. By eliminating *intI1*-harboring resistance plasmids in bacteria, we knock out both the class 1 integrase but also any, potentially unknown, ARGs in the cassette array and elsewhere on a given plasmid. Since *intI1* is often found on plasmids^24^, their elimination will not affect the viability of host bacteria, theoretically limiting the selective pressure for mutations that evade the CRISPR-Cas activity. In this study, we analyze the sequence conservation of *intI1* in the RefSeq database and investigate its distribution across plasmid taxonomic units (PTUs)^38^ in clinically relevant bacteria. We insert an *intI1*-targeting crispr RNA (crRNA) into the conjugative plasmid pNuc-cis encoding a TevCas9 dual nuclease^39^. We then show that transfer of this conjugative plasmid leads to curing of a resistance plasmid and consequent re-sensitization of antibiotic-resistant bacteria. This study shows the potential for targeting conserved resistance markers, such as *intI1*, for re-sensitization of antibiotic-resistant bacteria.

## Methods

### Assessment of plasmids containing class 1 integrons in RefSeq200

The 23,310 plasmids available in the NCBI RefSeq200 database as of May 20, 2020 were acquired. The database was manually curated, eliminating sequences that corresponded to partial plasmid DNA sequences, bacterial/archaeal chromosomes, or PacBio internal control sequences. This resulted in a curated dataset of 19,017 sequences. Plasmids were classified into PTUs using COPLA^40^. The host range of a PTU was defined according to Redondo-Salvo et al.^38^, categorizing them into six levels (I-VI) based on their observed distribution across taxonomic levels. These PTU host range levels are arranged from narrow to broad repertoires and are defined as I) within the same taxonomic species, II) within the same genus, III) within the same family, IV) within the same order, V) within the same class, and VI) within the same phylum.

A BLASTn^41^ search (with an E-value threshold of ≤10E−10 and minimum identity and query coverage of 90%) was conducted to identify class 1 integrons within the 19,017 plasmids, using the *intI1* gene from plasmid R388 as query (acc. no. FAA00063.1). Antibiotic resistance genes situated on plasmids with *intI1* were identified using BLASTn against the CARD^42^ database. Hits with ≥80% nucleotide identity, ≥80% sequence coverage, as used elsewhere^23^, and an E-value threshold of ≤10−5 were considered indicative of ARGs within class 1 integron-containing plasmids.

### Sequence conservation analysis of *intI1*

To analyze the sequence conservation of the *intI1* integrase gene of class 1 integrons, 2,628 *intI1* genes were identified from ESKAPEE organisms (*Enterococcus faecium, Staphylococcus aureus, Klebsiella pneumoniae, Acinetobacter baumannii, Pseudomonas aeruginosa, Enterobacter* spp., and *Escherichia coli*), using BLASTn against the NCBI RefSeq genome database. The search was performed in December 2023, and all hits with an E-value lower than 10E-10 and identity and query coverage above 90% were included. Sequences were oriented to the same strand and multiple sequence alignment was performed in Qiagen’s CLC Main Workbench 22.0.2 with default parameters.

### Design of *intI1*-targeting crRNA for targeted cleavage by Cas9

The IncP RK2-based plasmid pNuc-cis^43^ was selected as the conjugative plasmid to be used for cloning and re-sensitization. pNuc-cis was a gift from David Edgell (Addgene plasmid # 202796; http://n2t.net/addgene:202796; RRID:Addgene_202796). Plasmid pNuc-cis encodes a TevCas9 dual nuclease, combining the I-TevI nuclease with Cas9 to enhance target specificity, under the control of an arabinose-inducible P_*araBAD*_ promoter^39,43,44^. To ensure optimal cutting efficiency, the target site on the R388 *intI1* gene requires a 5’-NGG-3’ PAM sequence 3-4 nucleotides downstream from the Cas9 target site and a TevI cleavage motif (5’-CNNNG-3’) 15-18 nucleotides upstream^43^. A suitable target site within *intI1* was identified, based on sequence conservation, and oligonucleotides for the target site were synthesized with 4 bp overhangs complementary to the BsaI cut site on pNuc-cis (Oligo 1: 5’-CACGCTTGCCGTAGAAGAACAGCAG-3’, Oligo 2: 5’-AAAACTGCTGTTCTTCTACGGCAAG-3’; purchased from Integrated DNA Technologies, Inc. (IDT)). These served as *intI1* crRNA for the TevCas9 dual nuclease system.

### Inserting *intI1*-targetting crRNA into conjugative plasmid pNuc-Cis

Oligonucleotides (2 μl of 100 μM each) were phosphorylated using 2 μl of T4 Polynucleotide Kinase (catalogue no. M0201S; New England Biolabs, MA, USA), 5 μl T4 Polynucleotide Kinase Reaction Buffer (catalogue no. B0201S; New England Biolabs, MA, USA), and 2 μl of 25 mM ATP. Reaction volume was adjusted to 50 μl with ddH_2_O and incubated at 37°C for 30 minutes followed by heat inactivation at 75°C for 10 minutes. Annealing of the phosphorylated oligos was performed by adding 2.5 μl of 1M NaCl and heating to 95°C for 5 minutes followed by slow cooling to room temperature. For cloning, pNuc-cis was digested with BsaI (1 μL per 1-2 μg DNA; catalogue no. R0535; New England Biolabs, MA, USA) and Fast Digest Buffer (5 μL per 1-2 μg DNA) (catalogue no. B64; Thermo Fisher Scientific, MA, USA) at 37°C overnight. Digestion was verified by gel electrophoresis on a 1% agarose gel, comparing digested and undigested plasmid samples. Digested vector was purified via salt and ethanol precipitation, incubated at −20°C overnight, followed by centrifugation and ethanol washes. Ligation of the digested pNuc-cis vector and *intI1* crRNA insert was carried out at a 3:1 insert-to-vector ratio using T4 DNA Ligase (catalogue no. M0202S; New England Biolabs, MA, USA). Incubation was done at room temperature for 10 minutes, followed by 2 minutes at 37°C and inactivation at 65°C. A control ligation without insert was included to test for self-ligation of pNuc-cis. Ligates were quantified by Qubit 2.0 (Thermo Fisher Scientific, MA, USA).

### Transformation

To confirm correct cloning of pNuc-cis (pNuc-cis-crRNA_*intI1*_), the cloned DNA was transformed into electrocompetent *E. coli* ElectroMAX DH10B (Thermo Fisher Scientific, MA, USA) using five samples: a pNuc-cis positive control, a water negative control, a pNuc-cis digest, a ligated pNuc-cis without insert, and the cloned pNuc-cis-crRNA_*intI1*_. For each transformation, 50-100 ng/μL of DNA was added (water for the negative control) to 20 μL DH10B cells. Samples were electroporated at 25 μF, 200 Ω, and 1.66 kV using a Bio-Rad Gene Pulser II (Bio-Rad Laboratories, CA, USA). After electroporation, 950 μL S.O.C. medium was added, and samples were incubated at 37°C for 1 hour to express the Gm resistance phenotype. Each sample (90 μL) was plated on LB and LB with gentamicin (Gm, 25 μg/mL) plates to select for pNuc-cis transformants, with the remainder pelleted, resuspended in S.O.C. medium, and also plated on LB+Gm plates. Plates were incubated overnight at 37°C and colony growth was assessed the following day.

Fifty colonies from LB+Gm plate (*E. coli* transformed with cloned pNuc-cis-crRNA_*intI1*_) were selected for sequencing. Overnight cultures were grown in LB medium with 25 μg/mL gentamicin at 37°C. To minimize workload and cost of plasmid extraction, seventeen pools of clones were created, each containing 3 clones (3.3 mL per clone, 10 mL total per pool). Plasmid extraction was performed using the Plasmid Mini AX kit (A&A Biotechnology, Gdańsk, Poland). DNA concentrations were measured with the Qubit BR Assay (Thermo Fisher Scientific, MA, USA), and quality was assessed using a Nanodrop ND-1000 spectrophotometer (Thermo Fisher Scientific, MA, USA). DNA concentration was adjusted to 5 ng/μL in 10 μL molecular-grade water for Nanopore sequencing. Verification of clones was performed by sequencing with the Oxford Nanopore Technologies platform. The Rapid Barcoding Sequencing kit (SQK-RBK114.24) was used to build sequencing libraries and sequenced on a MinION R10.4.1 flow cell. Reads were basecalled with Guppy v6.5.7+ca6d6af (Oxford Nanopore Technologies) using the “dna_r10.4.1_e8.2_400bps_sup” model. Adapter sequences were removed using Porechop^45^ v0.2.4, followed by read filtration to achieve approximately 1 million base pairs per pool (100x coverage) with NanoFilt v2.8.0^46^. Flye v2.9.2-b1786^47^ was used to assemble basecalled reads. Mapped sequences of cloned pNuc-cis-crRNA_*intI1*_ were investigated for inserted *intI1* crRNA using CLC Genomics Workbench (Qiagen, Aarhus, Denmark).

### Conjugative transfer of pNuc-cis to antibiotic-resistant bacteria

To investigate the potential of re-sensitizing antibiotic-resistant bacteria by targeting class 1 integrons with a nuclease attack, we transferred the cloned pNuc-cis-crRNA_*intI1*_ by conjugation to recipient *E. coli* MG1655 with the IncW plasmid R388 that harbors *intI1*.

Overnight cultures of donor and recipient bacteria were grown in LB medium with antibiotics at 37°C (see Table 1). The next day, 0.2 μm pore size cellulose acetate filters (Sartorius, Göttingen, Germany) were placed on antibiotic-free LB agar plates. Each conjugation setup used one filter, plus an additional filter for each strain as a control. Cultures (1 mL each) were centrifuged, washed with LB, and resuspended in 500 μL LB. Equal volumes (50 μL) of resuspended donor and recipient cultures were placed onto filters. Plates were incubated at 37°C for 4 hours, after which filters were vortexed in 1 mL sterile PBS. A volume of 100 μL bacterial suspension was plated at different dilutions (10E^-0^ to 10E^-8^) on selective LB agar plates with gentamicin (Gm, 25 μg/mL; selective for donor and plasmid recipient bacteria), trimethoprim (Tm, 32 μg/mL; selective for target bacteria), or both antibiotics (selective for target bacteria that receive pNuc-cis but retain the target plasmid). Plates were supplemented with 0.2% glucose or 0.2% arabinose to regulate TevCas9 expression via P_*araBAD*_ in transconjugants. A control mating assay using crRNA_*intI1*_-lacking pNuc-cis donor strains was also performed. Plates were incubated overnight at 37°C, and colonies were counted.

**Table 1:** List of donor and target strain used for filter mating assay. The abbreviations for antibiotics are: gentamicin (Gm), nalidixic acid (Nx), trimethoprim (Tm). Strain DH5α carries ARG for nalidixic acid on the chromosome, while the other resistances are attributable to plasmids.

|  |
| --- |
| **Donor** |
| *E. coli* DH10B pNuc-cis-crRNA<sub>int1</sub> (Gm<sup>R</sup>) |
| *E. coli* DH5α pNuc-cis (Nx<sup>R</sup>, Gm<sup>R</sup>) |
| **Target bacteria** |
| *E. coli* MG1655 R388 (Tm<sup>R</sup>) |

## Results and Discussion

### Distribution of *IntI1* across PTUs and bacterial genera

To investigate the range of *intI1*-harboring resistance plasmids that can be targeted by the Cas9 system, we analyzed the prevalence of *intI1* within the RefSeq200 database. To enhance our understanding of this distribution, we employed the classification of Plasmid Taxonomic Units (PTUs). We identified *intI1* in 1,287 plasmids (Supplemental Table 1). Among these, 1,129 plasmids were classified into PTUs, while 158 remained unclassified. In total, 78 distinct PTUs contained *intI1*. Six PTUs (cPTU-939_2, cPTU-609_0, PTU-E55, cPTU-1187_7, cPTU-880_2, and PTU-E45) carried *intI1* in all plasmid members (Fig. 1). Our analysis reveals a diverse distribution of *intI1* within PTUs, underscoring its prevalence across different plasmid types (Fig. 1).

**Figure 1:**
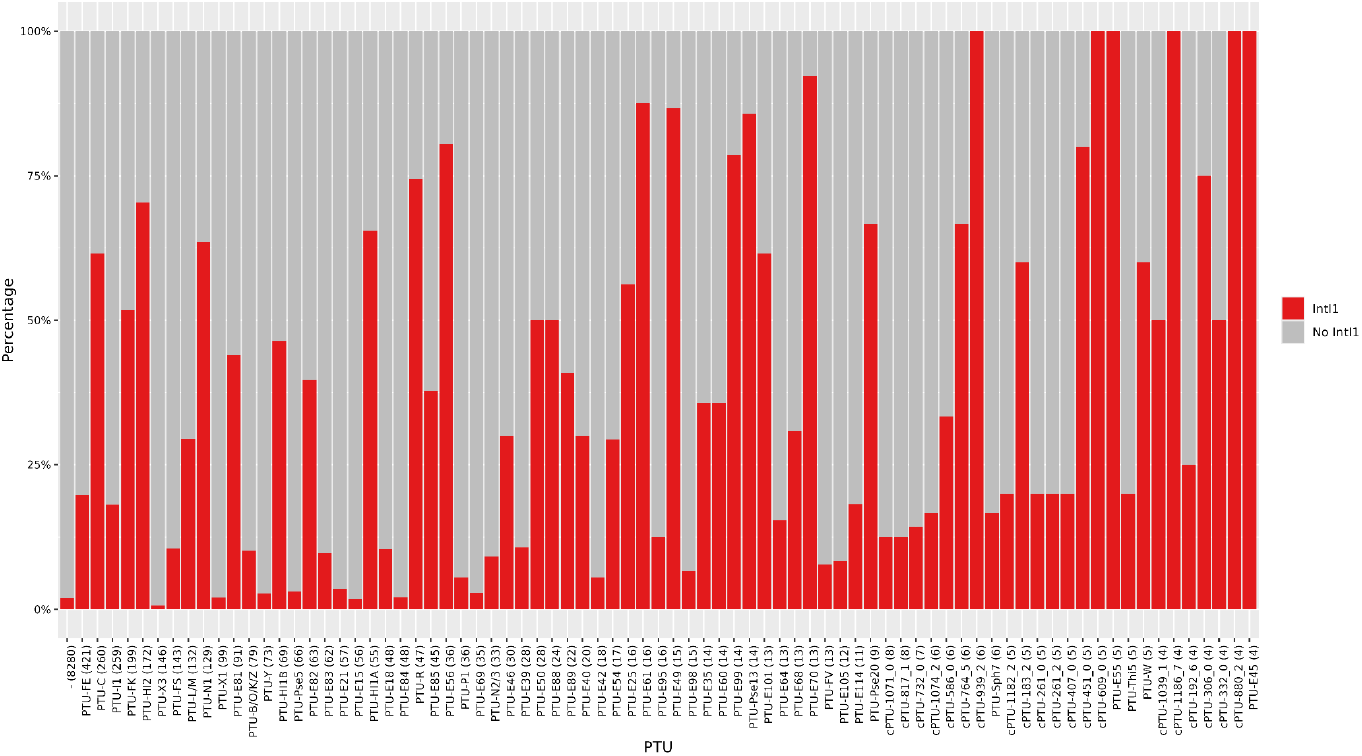
Proportion of intI1-Positive Plasmids in intI1-Containing PTUs. The barplot illustrates the different PTUs from RefSeq200 dataset that contain at least one intI1. PTUs are ordered on the X-axis according to the abundance of their members (indicated in parentheses). Each bar is color-coded to represent the percentage of plasmids within each PTU that harbor intI1 (red) and those that do not (grey). The first bar corresponds to plasmids not assigned to any PTU.

The host range assessment of *intI1*-containing PTUs (Fig. 2) was investigated. A large number of *intI1*-harboring PTUs (55 out of 78) have a host range III or higher, suggesting their ability to colonize bacteria from at least different genera (Fig. 2A).

**Figure 2:**
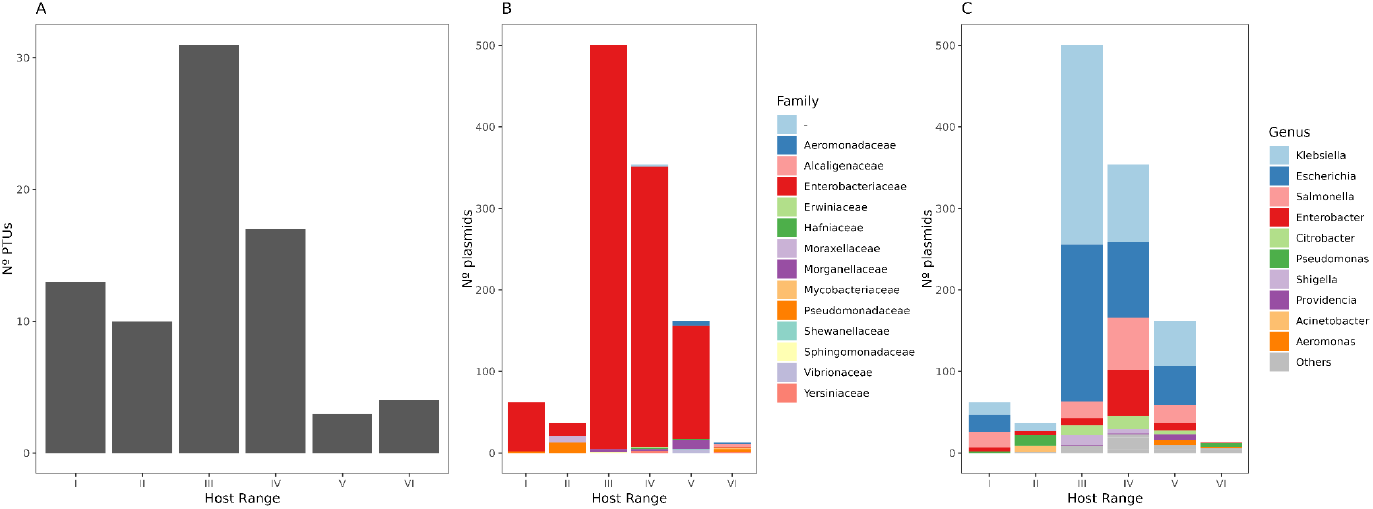
Host range of PTUs containing intI1. (A) Number of PTUs in each host range category. (B) Number of plasmids per host range, classified by family. Each bar represents the total count of plasmids in each host range, with colors indicative of the taxonomic families they belong to. (C) Number of plasmids per host range, classified by genus. Each bar represents the total count of plasmids in each host range, with colors indicative of the genera they belong to. Host ranges are I) within-species, II) within-genus, III) within-family, IV) within-order, V) within-class, and VI) within-phylum.

By delving into the prevalence of these plasmids across various bacterial families (Fig. 2B) and genera (Fig. 2C), we acquire insights into specific taxonomic groups harboring PTUs with *intI1* within the RefSeq200 dataset. The foremost family observed is *Enterobacteriaceae*, a result potentially influenced by the bias towards this family in the RefSeq200 dataset. Concerning genera, prominent ones include *Escherichia, Klebsiella*, and *Salmonella*. Notably, *Klebsiella* and *Escherichia* are part of the ESKAPEE group, known for their multidrug resistance and association with infections. While *Salmonella* is not categorized within the ESKAPEE group, it remains highly recognized for its pathogenicity and resistance traits^48^.

All plasmids that contain *intI1* in RefSeq200 contain ARGs (Supplemental Table 2). Figure 3 presents a heatmap depicting the number of plasmids per PTU that contain *intI1* and ARGs categorized by drug classes. This analysis sheds light on the potential drug classes that could be targeted when using a class 1 integron-targeting Cas9 system. PTU-C, PTU-HI2, PTU-FK, PTU-FE, PTU-HI1A, PTU-HI1B, and unclassified PTUs (-) have a high number of plasmids containing both *intI1* and ARGs against a wide range of antibiotics, with 290 distinct ARGs identified (Supplemental Table 2). This shows that targeting *intI1* could lead to re-sensitization towards multiple classes of antibiotics, offering a potential strategy to address MDR. The median number of ARGs per integron was 10 with a maximum of 38.

**Table 2:** Conjugation frequency and plasmid elimination efficiency.

| Sample | Conjugation frequency | Plasmid elimination efficiency |
| --- | --- | --- |
| *E. coli* pNuc-cis-crRNA<sub>int1</sub> + *E. coli* R388 | 5E-04 | 99.1 |
| *E. coli* pNuc-cis + *E. coli* R388 | 3E-03 | 28.2 |

**Figure 3:**
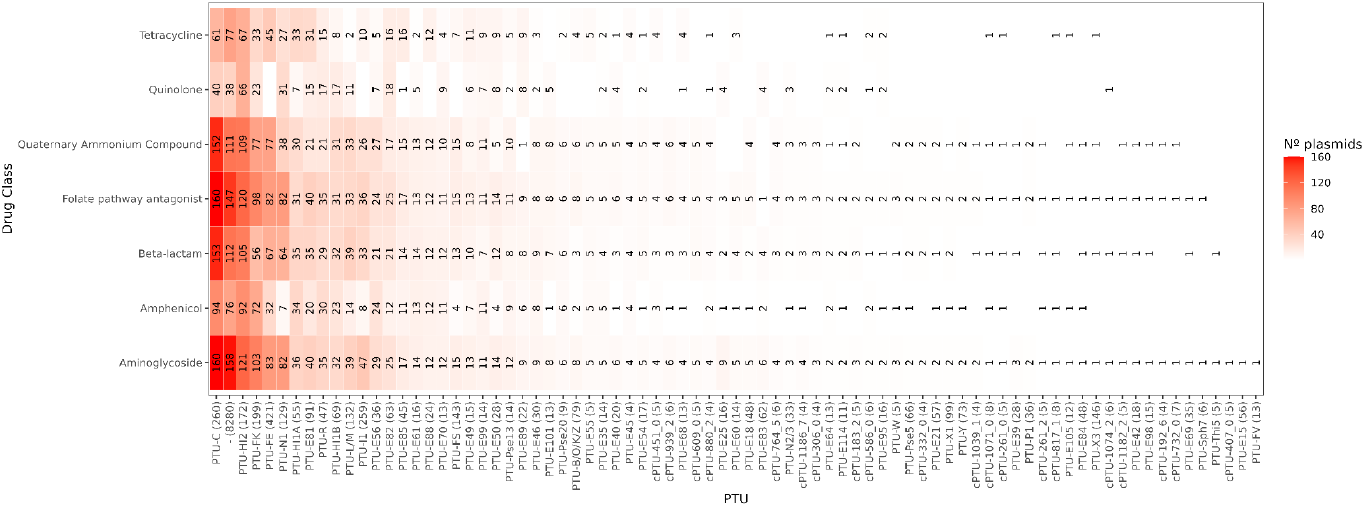
Distribution of drug classes in PTUs containing intI1. The X-axis represents the PTUs (with the total number of plasmids in RefSeq200 shown in parentheses). The Y-axis displays the drug classes. Each cell indicates the number of plasmids within each PTU that contain the corresponding drug class.

The top 10 most abundant ARGs on *intI1*-harboring plasmids are the aminoglycoside resistance-associated *AAC(6’)-Ib7* (1,188 plasmids), *ANT(3’’)-IIa* (425), *APH(6)-Id* (386), *APH(3’’)-Ib* (383), and *aadA2* (375), sulfonamide resistance-associated *sul1* (906) and *sul2* (432), antiseptic-resistance associated *qacEΔ1* (902), beta-lactamase *TEM-181* (599), and tetracycline efflux pump-encoding *tetA* (361). Other clinically significant ARGs are likewise residing on *intI1*-harboring plasmids, e.g. CTX-M, OXA, NDM beta-lactamases, and MCR variants for colistin resistance (Supplemental Table 2).

### The sequence of *intI1* is highly conserved

Alignment of the nucleotide sequence conservation analysis of 2,628 *intI1* genes from ESKAPEE organisms (BLASTn using *intI1* from R388 as a query against RefSeq genome database; E-value < 10E-10, >90% ID and query coverage) revealed an average sequence conservation of 99.997% per position (Fig. 4), which is in line with previous findings^24^. For ARGs that have previously been used as targets for CRISPR-Cas-based re-sensitization^4,30,31,49^, multiple subgroups exist based on sequence variants, making it near impossible to design crRNAs that broadly target e.g., most beta-lactamases. As an example, the TEM family of beta-lactamases currently comprises 250 variants (sub-terms) in the CARD^32^ database. The high sequence conservation of *intI1* facilitates the design of a crRNA that targets a universally conserved region (Fig. 4).

**Figure 4:**
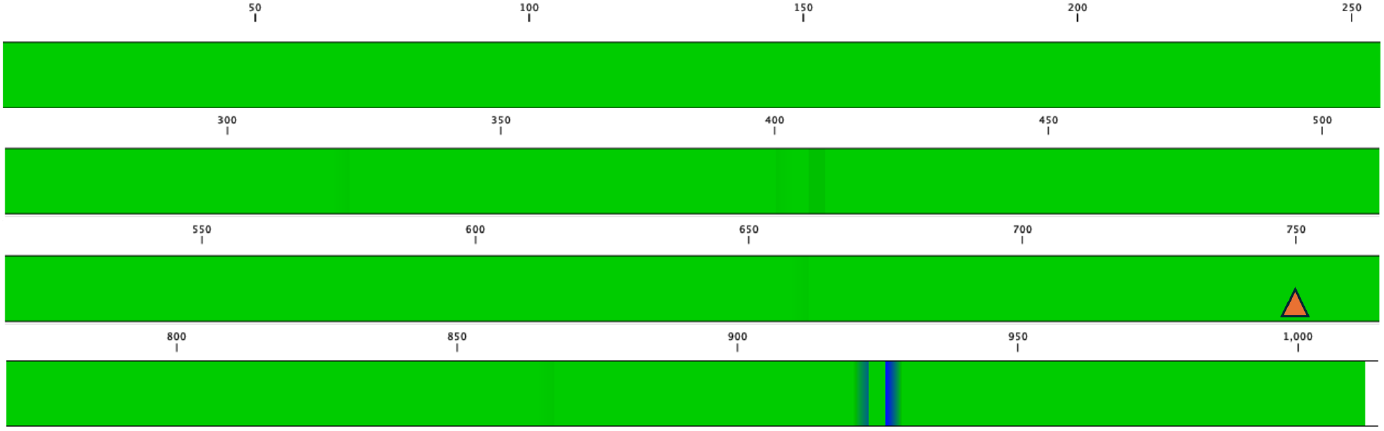
Nucleotide sequence conservation of class 1 integron integrase (intI1). A complete intI1 sequence (1,014 bp) from R388 was searched against the NCBI RefSeq genome database with BLASTn. Only ESKAPEE organisms were included to focus on a group of clinically relevant bacterial pathogens. 2,628 sequences were found, and sequences below 1,000 bp were then excluded (≈ 20 sequences) to avoid sequences with frameshift mutations resulting in possibly early stop codons and thereby nonfunctional proteins. Multiple sequence alignment was performed in CLC Genomics Workbench v. 24. Regions with 100% sequence conservation are shown in bright green, whereas dark blue indicates <99% conservation for a given region. Darker shades of green indicate between 99% and 100% conservation. A red triangle marks the selected target site for crRNA.

### Re-sensitization of antibiotic resistant *E. coli* by plasmid curing

After choosing an *intI1* target sequence (Fig. 4), the corresponding crRNA was cloned into pNuc-cis to re-sensitize antibiotic resistant bacteria by eliminating resistance plasmids. Successful insertion of *intI1*-targeting crRNA into pNuc-cis^43^ was validated with Oxford Nanopore sequencing. To demonstrate re-sensitization of antibiotic-resistant bacteria, the cloned pNuc-cis-crRNA_*intI1*_ plasmid was transferred via conjugation to *E. coli* MG1655 carrying class 1 integron on resistance plasmid R388. A conjugation frequency was calculated for each sample as transconjugants divided by target bacteria plus transconjugants:

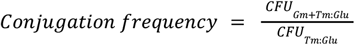

Conjugative delivery of pNuc-cis-crRNA_*intI1*_ to *E. coli* carrying R388 resulted in reduced transconjugant numbers (Gm^R^, Tm^R^) under TevCas9 expression induced by arabinose (Supplemental Table 3). This shows successful cleavage of *intI1* and elimination of plasmid R388. The co-existence of pNuc-cis-crRNA_*intI1*_ and R388 in a few remaining transconjugants, despite TevCas9 expression, may result from mutational inactivation of the system^50^, possibly through mutations in the target site or the CRISPR-Cas array. Conjugative delivery of pNuc-cis (without crRNA_*intI1*_ insertion) to *E. coli* R388 did not reduce transconjugants under induced TevCas9 conditions (arabinose) compared to repressed conditions (glucose), as expected. Plasmid elimination efficiency was calculated as a percentage of transconjugants in which target plasmid R388 is removed:

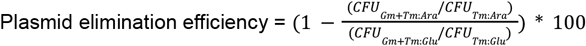

These efficiencies were calculated for each sample on the least diluted plates that allowed for CFU counting on the transconjugant Gm+Tm plates, i.e. between undiluted and 10E-2 (Supplemental Table 3). This plasmid elimination efficiency was 99.1% for pNuc-cis-crRNA_*intI1*_, compared to a background exclusion of 28.2% for pNuc-cis (Table 2). Cutting of *intI1* and R388 clearance may still occur under repressive TevCas9 conditions, likely due to leaky expression from the P_*araBAD*_ promoter (0.2% glucose), as previously reported. A reduction in conjugation frequency was observed for pNuc-cis-crRNA_*intI1*_ relative to pNuc-cis (Table 2).

When selecting for donor bacteria carrying either pNuc-cis-crRNA_*intI1*_ or pNuc-cis (Gm^R^) under arabinose conditions, both very small colonies and a few larger, countable colonies were observed (not shown). Hamilton et al. (2019)^43^ reported cellular toxicity resulting from leaky TevCas9 expression, which was further intensified by arabinose induction. This may account for the tiny colonies observed on Gm plates, but requires further investigation. During testing of pNuc-cis against other target plasmids, we found an instance of homologous recombination between the *pecM* gene on pNuc-cis and an identical gene on the target plasmid (pKJK5^51^) that led to loss of the cas9-encoding gene on pNuc-cis (results not shown). Future efforts should seek to further minimize conjugative plasmid vectors (e.g. pX1.0^52^) for delivery of Cas-based antimicrobials to reduce the risk of such recombination events. In designing plasmid vectors, codon optimization can be further applied to tailor vectors to specific bacterial targets. Alternative non-plasmid delivery vectors should also be further explored, as discussed recently^27^.

### Concluding remarks and future perspectives

The rapid rise and spread of antibiotic-resistant pathogens, especially due to MDR facilitated by HGT of MGEs and integrons, challenges the management of infectious diseases and calls for innovative solutions. The high specificity of the CRISPR-Cas system and its ability to be reprogrammed make it a tool to target resistance elements. Other studies have successfully used nucleases to re-sensitize resistant bacteria to β-lactams^4,30,31^, erythromycin^53^, tetracycline^53^, gentamicin^54^, colistin^55^, rifampicin^56^, and fosfomycin^57^. Furthermore, an alternative procedure has also been explored, where the conjugative acquisition of ARG plasmids were blocked^54^. While these studies demonstrate efforts in employing CRISPR-Cas technology against ARGs, they focus on single targets. Pursuing an individual or a limited number of ARGs at a time represents an indefinite effort, given the thousands of known ARGs^58^ and a similar high amount of different ARG-encoding plasmids^59^. This challenge is further complicated by the continuous emergence of new mutations in ARGs, which are selected under antibiotic exposure. In addition, defining what even qualifies as a significant resistance gene among clinically relevant pathogens also becomes important. Despite the option of employing multiple crRNAs in a single vector to mitigate mutational issues and variants, we maintain that the highly conserved *intI1* is the best target for nuclease-based re-sensitization.

In this study, we find 290 distinct ARGs co-occurring on *intI1*-harboring plasmids, showing the potential for re-sensitization towards a very broad range of antibiotics, by plasmid elimination through targeting the more universal *intI1*. By focusing on disrupting the spread of class 1 integron-containing plasmids, it may be possible to limit the transfer of resistance genes among bacterial populations of different hosts and mitigate the emergence and spread of multidrug-resistant pathogens. Despite promising results, future studies should aim to develop minimal conjugative plasmids for delivery, ensuring no sequence similarity to plasmids in target bacteria to avoid deletion of nuclease-encoding genes by homologous recombination. Other delivery vectors, such as nanoparticles, should also be further explored.

## References

1. WHO. No time to wait: Securing the future from drug-resistant infections. Artforum Int. (2019).

2. Tacconelli, E. et al. Discovery, research, and development of new antibiotics: the WHO priority list of antibiotic-resistant bacteria and tuberculosis. Lancet Infect. Dis. 18, 318–327 (2018).

3. Årdal, C. et al. To the G20: incentivising antibacterial research and development. Lancet Infect. Dis. 17, 799–801 (2017).

4. Tagliaferri, T. L. et al. Exploring the Potential of CRISPR-Cas9 Under Challenging Conditions: Facing High-Copy Plasmids and Counteracting Beta-Lactam Resistance in Clinical Strains of Enterobacteriaceae. Front. Microbiol. 11, 578 (2020).

5. Gholizadeh, P. et al. How CRISPR-Cas System Could Be Used to Combat Antimicrobial Resistance. Infect. Drug Resist. Volume 13, 1111–1121 (2020).

6. Tao, S., Chen, H., Li, N. & Liang, W. The Application of the CRISPR-Cas System in Antibiotic Resistance. Infect. Drug Resist. Volume 15, 4155–4168 (2022).

7. Zhang, H. et al. Resensitizing tigecycline- and colistin-resistant Escherichia coli using an engineered conjugative CRISPR/Cas9 system. Microbiol. Spectr. 12, e0388423 (2024).

8. Rafiq, M. S. et al. CRISPR-Cas System: A New Dawn to Combat Antibiotic Resistance. BioDrugs Clin. Immunother. Biopharm. Gene Ther. 38, 387–404 (2024).

9. Arnold, B. J.Huang, I.-T. & Hanage, W. P. Horizontal gene transfer and adaptive evolution in bacteria. Nat. Rev. Microbiol. 20, 206–218 (2022).

10. Shetty, V. P., Akshay, S. D., Rai, P. & Deekshit, V. K. Integrons as the potential targets for combating multidrug resistance in Enterobacteriaceae using CRISPR-Cas9 technique. J. Appl. Microbiol. 134, lxad137 (2023).

11. Bhat, B. A. et al. Integrons in the development of antimicrobial resistance: critical review and perspectives. Front. Microbiol. 14, (2023).

12. Ghaly, T. M. & Gillings, M. R. New perspectives on mobile genetic elements: a paradigm shift for managing the antibiotic resistance crisis. Philos. Trans. R. Soc. Lond. B. Biol. Sci. 377, 20200462 (2022).

13. Razavi, M., Kristiansson, E.Flach, C.-F. & Larsson, D. G. J. The Association between Insertion Sequences and Antibiotic Resistance Genes. mSphere 5, 10.1128/msphere.00418-20 (2020).

14. Gillings, M. R. Integrons: Past, Present, and Future. Microbiol. Mol. Biol. Rev. 78, 257 LP – 277 (2014).

15. Castañeda-Barba, S., Top, E. M. & Stalder, T. Plasmids, a molecular cornerstone of antimicrobial resistance in the One Health era. Nat. Rev. Microbiol. 22, 18–32 (2024).

16. Martínez, J. L., Coque, T. M. & Baquero, F. What is a resistance gene? Ranking risk in resistomes. Nat. Rev. Microbiol. 13, 116–123 (2015).

17. Partridge, S. R., Kwong, S. M., Firth, N. & Jensen, S. O. Mobile genetic elements associated with antimicrobial resistance. Clin. Microbiol. Rev. 31, 1–61 (2018).

18. Kamruzzaman, M. et al. Relative strengths of promoters provided by common mobile genetic elements associated with resistance gene expression in Gram-negative bacteria. Antimicrob. Agents Chemother. 59, 5088–5091 (2015).

19. Halaji, M. et al. The Global Prevalence of Class 1 Integron and Associated Antibiotic Resistance in Escherichia coli from Patients with Urinary Tract Infections, a Systematic Review and Meta-Analysis. Microb. Drug Resist. Larchmt. N 26, 1208–1218 (2020).

20. Gillings, M. et al. The evolution of class 1 integrons and the rise of antibiotic resistance. J. Bacteriol. 190, 5095–5100 (2008).

21. Ghaly, T. M. et al. The Natural History of Integrons. Microorganisms 9, 2212 (2021).

22. Hipólito, A., García-Pastor, L., Vergara, E., Jové, T. & Escudero, J. A. Profile and resistance levels of 136 integron resistance genes. Npj Antimicrob. Resist. 1, 1–12 (2023).

23. Nielsen, T. K., Browne, P. D. & Hansen, L. H. Antibiotic resistance genes are differentially mobilized according to resistance mechanism. GigaScience 11, giac072 (2022).

24. Zhang, A. N. et al. Conserved phylogenetic distribution and limited antibiotic resistance of class 1 integrons revealed by assessing the bacterial genome and plasmid collection. Microbiome 6, 130 (2018).

25. Baharoglu, Z., Bikard, D. & Mazel, D. Conjugative DNA Transfer Induces the Bacterial SOS Response and Promotes Antibiotic Resistance Development through Integron Activation. PLOS Genet. 6, e1001165 (2010).

26. Gillings, M. R., Krishnan, S., Worden, P. J. & Hardwick, S. A. Recovery of diverse genes for class 1 integron-integrases from environmental DNA samples. FEMS Microbiol. Lett. 287, 56–62 (2008).

27. Mayorga-Ramos, A., Zúñiga-Miranda, J., Carrera-Pacheco, S. E., Barba-Ostria, C. & Guamán, L. P. CRISPR-Cas-Based Antimicrobials: Design, Challenges, and Bacterial Mechanisms of Resistance. ACS Infect. Dis. 9, 1283–1302 (2023).

28. Duan, C., Cao, H.Zhang, L.-H. & Xu, Z. Harnessing the CRISPR-Cas Systems to Combat Antimicrobial Resistance. Front. Microbiol. 12, 716064 (2021).

29. Benz, F. et al. Type IV-A3 CRISPR-Cas systems drive inter-plasmid conflicts by acquiring spacers in trans. Cell Host Microbe 32, 875-886.e9 (2024).

30. Hao, M. et al. CRISPR-Cas9-Mediated Carbapenemase Gene and Plasmid Curing in Carbapenem-Resistant Enterobacteriaceae. Antimicrob. Agents Chemother. 64, e00843–20 (2020).

31. Kim, J.-S. et al. CRISPR/Cas9-Mediated Re-Sensitization of Antibiotic-Resistant Escherichia coli Harboring Extended-Spectrum β-Lactamases. J. Microbiol. Biotechnol. 26, 394–401 (2016).

32. Alcock, B. P. et al. CARD 2020: Antibiotic resistome surveillance with the comprehensive antibiotic resistance database. Nucleic Acids Res. 48, D517–D525 (2020).

33. Hall, B. G. & Barlow, M. Evolution of the serine beta-lactamases: past, present and future. Drug Resist. Updat. Rev. Comment. Antimicrob. Anticancer Chemother. 7, 111–123 (2004).

34. Bush, K. Past and Present Perspectives on β-Lactamases. Antimicrob. Agents Chemother. 62, e01076 (2018).

35. Fröhlich, C., Chen, J. Z., Gholipour, S., Erdogan, A. N. & Tokuriki, N. Evolution of β-lactamases and enzyme promiscuity. Protein Eng. Des. Sel. 34, gzab013 (2021).

36. Henderson, T. A., Young, K. D., Denome, S. A. & Elf, P. K. AmpC and AmpH, proteins related to the class C β-lactamases, bind penicillin and contribute to the normal morphology of Escherichia coli. J. Bacteriol. 179, 6112–6121 (1997).

37. Brandt, C. et al. In silico serine β-lactamases analysis reveals a huge potential resistome in environmental and pathogenic species. Sci. Rep. 7, 43232 (2017).

38. Redondo-Salvo, S. et al. Pathways for horizontal gene transfer in bacteria revealed by a global map of their plasmids. Nat. Commun. 11, 3602 (2020).

39. Wolfs, J. M. et al. Biasing genome-editing events toward precise length deletions with an RNA-guided TevCas9 dual nuclease. Proc. Natl. Acad. Sci. 113, 14988–14993 (2016).

40. Redondo-Salvo, S. et al. COPLA, a taxonomic classifier of plasmids. BMC Bioinformatics 22, 390 (2021).

41. Altschul, S. F., Gish, W., Miller, W., Myers, E. W. & Lipman, D. J. Basic local alignment search tool. J. Mol. Biol. 215, 403–410 (1990).

42. McArthur, A. G. et al. The Comprehensive Antibiotic Resistance Database. Antimicrob. Agents Chemother. 57, 3348–3357 (2013).

43. Hamilton, T. A. et al. Efficient inter-species conjugative transfer of a CRISPR nuclease for targeted bacterial killing. Nat. Commun. 10, 4544 (2019).

44. Guha, T. & Edgell, D. Applications of Alternative Nucleases in the Age of CRISPR/Cas9. Int. J. Mol. Sci. 18, 2565 (2017).

45. Wick, R. R., Judd, L. M., Gorrie, C. L. & Holt, K. E. Completing bacterial genome assemblies with multiplex MinION sequencing. Microb. Genomics 3, e000132 (2017).

46. De Coster, W., D’Hert, S., Schultz, D. T., Cruts, M. & Van Broeckhoven, C. NanoPack: visualizing and processing long-read sequencing data. Bioinformatics 34, 2666–2669 (2018).

47. Kolmogorov, M., Yuan, J., Lin, Y. & Pevzner, P. A. Assembly of long, error-prone reads using repeat graphs. Nat. Biotechnol. 37, 540–546 (2019).

48. WHO bacterial priority pathogens list, 2024: Bacterial pathogens of public health importance to guide research, development and strategies to prevent and control antimicrobial resistance. https://www.who.int/publications/i/item/9789240093461.

49. Li, P. et al. Targeted Elimination of blaNDM-5 Gene in Escherichia coli by Conjugative CRISPR-Cas9 System. Infect. Drug Resist. 15, 1707 (2022).

50. Uribe, R. V. et al. Bacterial resistance to CRISPR-Cas antimicrobials. Sci. Rep. 11, 17267 (2021).

51. Bahl, M. I., Hansen, L. H., Goesmann, A. & Sørensen, S. J. The multiple antibiotic resistance IncP-1 plasmid pKJK5 isolated from a soil environment is phylogenetically divergent from members of the previously established α, β and δ sub-groups. Plasmid 58, 31–43 (2007).

52. Hansen, L. H. et al. Design and synthesis of a quintessential self-transmissible IncX1 plasmid, pX1.0. PloS One 6, e19912 (2011).

53. Rodrigues, M., McBride, S. W., Hullahalli, K., Palmer, K. L. & Duerkop, B. A. Conjugative Delivery of CRISPR-Cas9 for the Selective Depletion of Antibiotic-Resistant Enterococci. Antimicrob. Agents Chemother. 63, e01454–19 (2019).

54. Walker-Sünderhauf, D. et al. Removal of AMR plasmids using a mobile, broad host-range CRISPR-Cas9 delivery tool. Microbiology 169, 001334 (2023).

55. Wan, P. et al. Reversal of mcr-1-Mediated Colistin Resistance in Escherichia coli by CRISPR-Cas9 System. Infect. Drug Resist. 13, 1171 (2020).

56. Faulkner, V. et al. Re-sensitization of Mycobacterium smegmatis to Rifampicin Using CRISPR Interference Demonstrates Its Utility for the Study of Non-essential Drug Resistance Traits. Front. Microbiol. 11, 619427 (2020).

57. Walflor, H. S. M., Lucena, A. R. C., Tuon, F. F., Medeiros, L. C. S. & Faoro, H. Resensitization of Fosfomycin-Resistant Escherichia coli Using the CRISPR System. Int. J. Mol. Sci. 23, 9175 (2022).

58. Liu, B. & Pop, M. ARDB - Antibiotic resistance genes database. Nucleic Acids Res. 37, D443–447 (2009).

59. Orlek, A., Anjum, M. F., Mather, A. E., Stoesser, N. & Walker, A. S. Factors associated with plasmid antibiotic resistance gene carriage revealed using large-scale multivariable analysis. Sci. Rep. 13, 2500 (2023).

